# Metabolic effects and biotransformation of perfluorohexyloctane (F6H8) in human hepatocytes

**DOI:** 10.1101/2025.10.24.684326

**Authors:** Andi Alijagic, Jade Chaker, João Marcos G. Barbosa, Daniel Duberg, Victor Castro-Alves, Alex M. Dickens, Matej Orešič, Tuulia Hyötyläinen

**Affiliations:** Man-Technology-Environment (MTM) Research Centre, School of Science and Technology, Örebro University, SE-701 82 Örebro, Sweden; Inflammatory Response and Infection Susceptibility Centre (iRiSC), Örebro University, Örebro SE-701 82, Sweden; School of Medical Sciences, Faculty of Medicine and Health, Örebro University, SE-701 82 Örebro, Sweden; Turku Centre for Biotechnology, University of Turku and Åbo Akademi University, FI-20520 Turku, Finland; Department of Chemistry, University of Turku, Finland, FI-20500 Turku, Finland; Department of Life Technologies, University of Turku, FI-20014 Turku, Finland

**Author notes:** **Contact information:** Tuulia Hyötyläinen, School of Science and Technology, Örebro University, 70182 Örebro, Sweden.

**Keywords:** **Keywords**: HepaRG, lipidomics, liver metabolism, metabolomics, perfluorohexyloctane, PFAS

## Abstract

Perfluorohexyloctane (F6H8) is a semifluorinated alkane recently approved for ophthalmic treatment of dry eye disease. Although considered locally safe for topical use, its structural similarity to persistent per- and polyfluoroalkyl substances (PFAS) raises concerns about systemic accumulation and long-term toxicity. To investigate potential hepatic effects, we examined the metabolic impact of F6H8 exposure in human HepaRG hepatocytes across a broad concentration range representing short- and long-term exposure scenarios. Untargeted metabolic profiling by ultra-high-performance liquid chromatography–quadrupole time-of-flight mass spectrometry (UHPLC-QTOFMS) was performed on intracellular extracts and extracellular media. F6H8 induced pronounced, concentration-dependent metabolic alterations, many of which exhibited non-monotonic responses. Low concentrations primarily affected amino acid, fatty acid, and lipid metabolism, while central carbon metabolism was disrupted only at the highest exposures. Notably, a putative biotransformation product, perfluorohexyloctanoic acid, was detected, suggesting metabolic persistence and conversion to a PFAS-like structure. These findings indicate that F6H8 elicits broad metabolic reprogramming and may not be metabolically inert as previously assumed. Given its clinical use and structural similarity to persistent fluorochemicals, the results highlight the need for comprehensive, long-term safety assessment of F6H8 and related semifluorinated alkanes.

## Introduction

Perfluorohexyloctane (F6H8) is a semifluorinated alkane approved for ophthalmic use and increasingly incorporated into ophthalmic formulations for the treatment of dry eye disease, particularly those associated with meibomian gland dysfunction. Clinical studies have demonstrated its efficacy and short-term safety following topical administration,^1, 2^ however, its broader toxicological profile remains poorly characterized. Owing to its classification within the per- and polyfluoroalkyl substance (PFAS) class, F6H8 shares physicochemical features with other highly fluorinated compounds that are known to persist in biological systems and the environment, raising concerns regarding potential systemic accumulation and long-term health effects.

Compared to well-studied PFAS such as perfluorooctanoic acid (PFOA) and perfluorooctane sulfonate (PFOS), the metabolic fate, biotransformation potential, and potential cellular effects of F6H8 remain poorly understood. Existing toxicological data are limited to short-term clinical safety evaluations under ophthalmic exposure scenarios. Notably, structurally related semifluorinated alkanes with longer hydrocarbon tails have been used in industrial applications such as ski waxes—a practice now restricted or banned in many countries because of concerns over environmental persistence and bioaccumulation. Analyses of ski wax formulations have shown that semifluorinated alkanes commonly contain fluorinated chains of F6H16, F10H16, F12H16, F14H16 and F16H16.^3^

There is limited data on biological effects of F6H8 from *in vivo* studies. Reported nonclinical pharmacology and toxicology studies include two studies in rabbits and two in rats.^4^ The endpoints analysed were toxicity in two of the studies and embryofoetal developmental (EFD) toxicity in the other two. In a 26-week ocular toxicity study in rabbits, ocular instillation of F6H8 ophthalmic solution (427.8 mg/day, four times daily bilaterally for up to 26 weeks) was well tolerated and did not induce any ocular or systemic signs of toxicity. In another 28-day oral toxicity study in rats, daily oral administration of at doses up to 2000 mg/kg/ for 28 days were well tolerated and did not produce F6H8 related toxicity. In an EFD toxicity study in rats, daily oral administration of F6H8 at doses up to 2000 mg/kg/day to pregnant Wistar rats during the period of organogenesis was well tolerated with no toxicological effects on maternal or embryofoetal parameters.^4^ However, in an EFD toxicity study in rabbits, following daily oral administration of F6H8 (0, 250, 500 and 1000 mg/kg/day) to pregnant female New Zealand white (NZW) rabbits during the period of organogenesis, there were abortions in all treated groups, compared with no abortion in the control group.^4^ In animals treated with F6H8, normal weight gain from birth to adulthood was reduced in a dose-dependent manner. Reduced food consumption was also noted at a dose-related trend. There were higher incidences of reduced fecal output, soft faeces and/or absent urine in all groups during the dosing period compared with the control group. Consistent with the maternal toxicity, mean fetal weight was reduced in all treated groups compared with the concurrent control group. There was no increase in embryofoetal death nor test article-related delay in skeletal ossification in any treated group, however, there were more fetal malformations (external, visceral, and/or skeletal) in the low, mid and high dose treatment groups as compared with the control group. In a rat study using F6H8-based propofol, elevation of liver enzyme alanine-aminotransferase (ALT) after exposure to F6H8-based propofol was observed in absence of histological changes in the liver, in comparison with conventional propofol emulsion.^5^ Viability and proliferation of cultured human corneal endothelial cells and human retinal pigment epithelial cells were decreased after incubation with F6H8, which was likely due to high lipophilicity of F6H8 interactions with cellular lipoprotein membranes.^6^ In a recent zebrafish model study, F6H8 was shown to induce a hypoactive embryonic photomotor response, suggesting potential developmental neurotoxicity.^7^ Clinical studies in humans have been primarily focused on ocular endpoints and symptom relief, lacking investigation of systemic and long-term chronic exposure effects.

There is very limited data on adsorption, organ distribution, and excretion of F6H8. In a rabbit study, labelled F6H8 was administered into eyes of the rabbits, and the concentration of F6H8 was investigated in ocular tissues, blood and plasma using radioactivity by liquid Scintillation counting.^8^ Following a single dose, plasma concentrations increased to 973 ng/g at 4 hours and declined to 360 ng/g by 24 hours, suggesting potential accumulation with chronic dosing of multiple times daily. Compounds with similar structure to F6H8, namely C_6_F_13_C =CHC_10_H_21_ (F6H10E), has been investigated with regards to the organ distribution and organ retention time in rats.^9^ The F6H10E content in the liver peaked one day after administration (seven days for the spleen). At a dose of 3.6 g/kg body weight, the hepatic half-life of F6H10E was estimated at 25 ± 5 days.^9^ In addition, saturated alkanes, *i.e.*, compounds with structure otherwise identical, but with no fluorine atoms, has been shown to enrich in the liver, where they are metabolized into carboxylic acids with the same chain length.^10–12^ The liver concentrations were ca. two times higher than in the blood.

Taken together, toxicological and chronic exposure data on F6H8 remain scarce, particularly regarding its biological effects. Existing investigations have primarily focused on acute toxicity, which may be insufficient to reveal the consequences of long-term chronic exposures, as adverse health effects can take extended periods to develop. To date, no published studies have addressed the metabolic effects of F6H8. The US Food and Drug Administration (FDA) approval documents state that F6H8 is not metabolized by human liver microsomes, however, no supporting data or peer-reviewed studies have been published.

Related work has demonstrated that semifluorinated alkanes can interact with phospholipids and integrate into lipid bilayers.^13^ When co-dispersed with phospholipids, the lipophobic fluorinated have been shown to aggregate and form a phase-separated fluorinated core within the membrane.^14^ Thus, even if F6H8 itself were metabolically inert, it could still alter cellular metabolism by modifying membrane structure and function.

To address this critical knowledge gap, we investigated the metabolic effects of F6H8 exposure in human hepatocytes, given the liver’s central role in xenobiotic metabolism and systemic toxicity. Using comprehensive mass spectrometry (MS) based untargeted metabolic profiling, we examined how F6H8 influences cellular metabolism across a wide concentration range and assessed possible biotransformation of the compound. Both intracellular extracts and extracellular media were analyzed to provide an integrated view of exposure-related metabolic changes. Our findings offer foundational toxicological insight into this widely used but poorly understood compound and inform future risk assessment of F6H8 and related fluorinated substances.

## Materials and methods

### Chemicals

All solvents were HPLC grade or LC-MS grade, from Honeywell (Morris Plains, NJ, USA), Fisher Scientific (Waltham, MA, USA) or Sigma-Aldrich (St. Louis, MO, USA). MS grade ammonium acetate and reagent grade formic acid were also from Sigma-Aldrich (St. Louis, MO, USA). The lipid standards were from Avanti Polar Lipids Inc. (Alabaster, AL, USA). ^13^C-labeled PFAS internal standards (IS), ^13^C-labeled performance standards, and native calibration standards (perfluorocarboxylic acids [PFCAs] and perflurosulfonic acids [PFSAs]) were purchased from Wellington Laboratories (Guelph, Ontario, Canada). One native performance standard, 7H-dodecafluoroheptanoic acid, was purchased from ABCR (Karlsruhe, Germany). Cholic acid (CA), chenodeoxycholic acid (CDCA), deoxycholic acid (DCA), dehydrocholic Acid (DHCA), glycocjolic acid (GCA), glyco chenodeoxycholic acid (GCDCA), lithocholic acid (LCA), taurocholic acid (TCA), taurochenodeoxycholic acid (TCDCA), taurodeoxycholic acid TDCA, taurodehydrocholic acid (TDHCA), taurohyocholic acid (THCA), taurohyodeoxycholic acid (THDCA), taurolithocholic acid (TLCA), and tauroursocholic acid (TUDCA) were obtained from Sigma-Aldrich (St. Luis, MO, USA), hyodeoxycholic acid (HDCA), hyocholic acid (HCA), murocholic acids α, β and ω (αMCA, βMCA, ωMCA), 7-oxohyodeoxycholic acid (7-oxo-HDCA), 7-oxodeoxycholic acid (7-oxo-DCA), 12-oxo lithocholic acid (12-oxo-LCA), tauromurocholic acid), murocholic acids α, β and ω (TαMCA, TβMCA, TωMCA), glycodehydrocholic acid (GDHCA), glycohyocholic acid (GHCA), and glycohyodeoxycholic acid) GHDCA from Steraloids (Newport, RI, U.S.A), glycolithocholic acid (GLCA) and glycoursocholic acid (GUDCA) from Calbiochem (Gibbstown, NJ, U.S.A), and glycodeoxycholic acid (GDCA) and ursocholic acid (UDCA) from Fluka (Buchs, Switzerland). Internal standards CA-d4, LCA-d4, UDCA-d4, CDCA-d4, DCA-d4, GCA-d4, GLCA-d4, GUDCA-d4 and GCDCA-d4 were obtained from Qmx laboratories Ltd. (Essex, UK). For quality assurance (QA), standard reference material serum NIST SRM 1950 (for lipidomics and metabolomics) and 1957 (for PFAS and bile acids) was purchased from the National Institute of Standards and Technology (NIST) at the US Department of Commerce (Washington, DC, USA).

Perfluorohexyloctane was purchased from a local pharmacy. It was analyzed by nuclear magnetic resonance (NMR) to verify its purity. The main compound had both fluorine and protein peaks that aligned with previous literature values.^15^ However, there was smaller fluorine peaks observed (**Supplementary Figure 1**) suggesting that there is other fluorinated species contained in the eye drop solution. The NMR was run on a Bruker Advance III 500 system equipped with a smart probe. Both F19 and 1H NMR was performed to confirm the structure of the eyedrop solution.

### Cell culture maintenance

Undifferentiated HepaRG® human hepatic cell line (BIOPREDIC, France; isolated from the female donor), also considered a surrogate for primary human hepatocytes,^16^ was cultured in the basal hepatic cell medium supplemented with HepaRG® Growth Medium Supplement with antibiotics (BIOPREDIC, France). Cells were grown in T75 cell culture flasks until reaching 100% confluency (usually 3-4 days). Later, cells were maintained in the same culture flask for an additional 14 days in order to promote differentiation into hepatocyte-like and biliary-like cells under DMSO-free conditions.^17^ Complete cell medium was changed every 2-3 days. The cells were maintained at 37°C with 5% CO_2_ atmosphere.

### Cell seeding, exposure, and sample collection

On day 1, cells were trypsinized and seeded in 24-well culture plates (VWR, Sweden) at the high density of 10^6^ cells per well in the volume of 1000 µL and incubated for an additional 24 h. On day 2, cells were exposed to perfluorohexyloctane (F6H8) in a volume of 500 µL, using five different concentrations designed to simulate short- and long-term exposure scenarios: corresponding to 1-, 7-, 14-, 21-, and 28-day systemic exposure (D1, D7, D14, D21 and D28, respectively). The rationale for the selected concentrations was based on an estimated exposure model. We assumed that approximately 5% of the administered ophthalmic dose could reach hepatic tissue. As per clinical usage, eye drops are typically prescribed up to four times per day, with one drop per administration. Given that a single drop contains approximately 11 µL of solution, this results in a total daily dose of 44 µL. Applying the 5% systemic availability assumption, we estimated hepatic exposure and calculated equivalent concentrations for use in the *in vitro* model. After 24-h exposure, 300 µL of the cell culture supernatant was aspirated and transferred to a 96-deepwell microplate. The remaining cell media was removed and 300 µL of pre-warmed PBS was added on top of cells. Cells were gently detached by the cell scraper and transferred to a 96-deepwell microplate. Samples were stored at -80°C until extraction and analysis. For each cell line, experiments included 4 technical replicates, and 3 independent experiments/biological replicates, yielding in total 12 replicates for each exposure condition.

### Cell viability assessment

HepaRG cells were seeded at high density in 96-well black-walled microplates (Revvity) at 50,000 cells/well and incubated at 37 °C with 5% CO₂ for 24 h. Afterwards, the medium was removed by inverting the plates, and the cells were exposed to the same concentrations of perfluorohexyloctane as described in the section *Cell seeding, exposure, and sample collection*. After 24 h of exposure, the alamarBlue assay was performed as described previously.^18^ In brief, the cell medium was removed, and 10% alamarBlue™ HS Cell Viability Reagent (Invitrogen; Thermo Fisher Scientific, Eugene, OR, USA) in pre-warmed basal hepatic cell medium was added. The cells were incubated for 3 h at 37 °C. Fluorescence was then measured using a multi-mode microplate reader (FLUOstar Omega, BMG LABTECH, Ortenberg, Germany) at 540/590 nm excitation/emission. Viability is expressed as relative fold change compared to the negative (unexposed) control (**Supplementary Figure 2**).

### Analysis of lipids, polar and semipolar metabolites

All samples were randomized before sample preparation and analysis. The samples were analyzed using two parallel methods, one aimed at analysis of polar and semipolar metabolites and the second one for the analysis of molecular lipids (**Table 1**). The cell pellets were first homogenized by adding 150 µL 0.9% NaCl solution, after which the samples were vortex mixed and ultrasonicated for 3 minutes. Both analyses were done by an ultra-high-performance liquid chromatography quadrupole time-of-flight mass spectrometry (UHPLC-QTOFMS). In short, the UHPLC system used for lipidomic analyses was a 1290 Infinity II system from Agilent Technologies (Santa Clara, CA, USA. MassHunter B.06.01 software was used for data acquisition. For the analysis of polar and semipolar compounds, analyzed using an Xevo G3 QTOF system (Waters Corporation). Data acquisition was done with MassLynx software.

**Table1.** LC-MS conditions for the two methods used in the study.

| Conditions | Polar/semipolar compounds:<br>Method 1 | Lipidomics: Method 2 |
| --- | --- | --- |
| **Injection volume** | 2 µl | 1 µl |
| **Column** | C18 precolumn (Waters Corporation, Wexford, Ireland) and an inline filter, pore size 0,2 µm (Waters Corporation, Wexford, Ireland) + ACQUITY UPLC® BEH C18 column (2.1 mm × 100 mm, particle size 1.7 µm) by Waters (Milford, MA, USA) | C18 precolumn (Waters Corporation, Wexford, Ireland) and an inline filter, pore size 0,2 µm (Waters Corporation, Wexford, Ireland) + ACQUITY UPLC® BEH C18 column (2.1 mm × 100 mm, particle size 1.7 µm) by Waters (Milford, MA, USA) |
| **Mobile phases** | A H<sub>2</sub>O:MeOH (v/v 70:30) with 2 mM ammonium acetate<br>B MeOH with containing 2 mM ammonium acetate | A 10 mM ammonium acetate and 0.1% Formic Acid in H<sub>2</sub>O<br>B Acetonitrile:Isopropanol (v/v 1:1) with 0.1% Formic Acid and 10 mM ammonium acetate |
| **Gradient** | • 0-1.5 min: B was increased from 5% to 30%<br>• 1.5-4.5 min: B increased to 70%<br>• 4.5-7.5 min: B increased to 100% and held for 5.5 min.<br>• A post-time of 6 minutes | • 0-2 min: B was increased from 35% to 80%<br>• 2-7 min: B increased to 100%<br>• 7-14 min: B was held at 100%<br>• A post-time of 7 min |
| **Flow rate** | 0.4 mLmin<sup>-1</sup> | 0.4 mLmin<sup>-1</sup> |
| **MS conditions** | Dual ESI ionization source with capillary voltage 4.5 kV, nozzle voltage 1500 V, N<sub>2</sub> pressure in the nebulized was 21 psi and the N<sub>2</sub> flow rate and temperature as sheath gas was 11 Lmin<sup>-1</sup> and 379 °C, respectively. Drying gas flow was set to 10 Lmin<sup>-1</sup> and temperature to 150 °C. m/z range 100-1700 in negative ion mode. | Dual ESI ionization source with capillary voltage 3.64 kV, nozzle voltage 1500 V, N<sub>2</sub> pressure in the nebulized was 21 psi and the N<sub>2</sub> flow rate and temperature as sheath gas was 11 Lmin<sup>-1</sup> and 379 °C, respectively. Drying gas flow was set to 10 Lmin<sup>-1</sup> and temperature to 193 °C. m/z range 100-1700 in positive ion mode |

80 µL of cell homogenate or 80 µL cell media was extracted with 480 µL of monophasic MeOH:MTBE:IPA (20:15:15, v/v) containing the following internal standards: Valine-d8, Glutamic acid-d5, Succinic acid-d4, Heptadecanoic acid, Lactic acid-d3, Citric acid-d4. 3-Hydroxybutyric acid-d4, Arginine-d7, Tryptophan-d5, Glutamine-d5, CA-d4, CDCA-d4, CDCA-d4, GCA-d4, GCDCA-d4, GLCA-d4, GUDCA-d4, LCA-d4, TCA-d4, UDCA-d4, 1,2-diheptadecanoyl-sn-glycero-3-phosphoethanolamine (PE(17:0/17:0)), N-heptadecanoyl-D-erythro-sphingosylphosphorylcholine (SM(d18:1/17:0)), N-heptadecanoyl-D-erythro-sphingosine (Cer(d18:1/17:0)), 1,2-diheptadecanoyl-sn-glycero-3-phosphocholine (PC(17:0/17:0)), 1-heptadecanoyl-2-hydroxy-sn-glycero-3-phosphocholine (LPC(17:0)) and 1-palmitoyl-d31-2-oleoyl-sn-glycero-3-phosphocholine (PC(16:0/d31/18:1)), and triheptadecanoylglycerol (TG(17:0/17:0/17:0)). After vortex-mixing (30 s), the extract was kept on an ice plate (45 min) and was filtered using filtration plates (0.45 µm). For polar/semipolar metabolite analysis, a 100 µL-aliquot was transferred to a vial, evaporated in a speed-vac system, and reconstituted with 50 µL MeOH:H_2_O (7:3, v/v) for analysis. For lipidomics analysis, the extracted sample was directly anayzed with no additional step.

Quantitation was done using 6-point calibration (bile acids c=20-640 ng/mL, polar metabolites c=0.1 to 80 μg/mL). Quantification of other bile acids was done using the following compounds: Chenodeoxycholic acid (CDCA), Cholic acid (CA), Deoxycholic acid (DCA), Glycochenodeoxycholic acid (GCDCA), Glycocholic acid (GCA), Glycodehydrocholic acid (GDCA), Glycodeoxycholic acid (GDCA), Glycohyocholic acid (GHCA), Glycohyodeoxycholic acid (GHDCA), Glycolithocholic acid (GLCA), Glycoursodeoxycholic acid (GUDCA), Hyocholic acid (HCA), Hyodeoxycholic acid (HDCA), Litocholic acid (LCA), alpha-Muricholic acid (αMCA), Tauro-alpha-muricholic acid (T-α-MCA), Tauro-beta-muricholic acid(T-β-MCA), Taurochenodeoxycholic acid (TCDCA), Taurocholic acid (TCA), Taurodehydrocholic acid (THCA), Taurodeoxycholic acid (TDCA), Taurohyodeoxycholic acid (THDCA), Taurolithocholic acid (TLCA), Tauro-omega-muricholic acid (TωMCA) and Tauroursodeoxycholic acid (TDCA) and polar metabolites was done using alanine, citric acid, fumaric acid, glutamic acid, glycine, lactic acid, malic acid, 2-hydroxybutyric acid, 3-hydroxybutyric acid, linoleic acid, oleic acid, palmitic acid, stearic acid, cholesterol, fructose, glutamine, indole-3-propionic acid, isoleucine, leucine, proline, succinic acid, valine, asparagine, aspartic acid, arachidonic acid, glycerol-3-phosphate, lysine, methionine, ornithine, phenylalanine, serine and threonine. The curves had R values >0.98 for most of the compounds. For lipidomics, calibration curves using 1-hexadecyl-2-(9Z-octadecenoyl)-sn-glycero-3-phosphocholine (PC(16:0e/18:1(9Z))), 1-(1Z-octadecenyl)-2-(9Z-octadecenoyl)-sn-glycero-3-phosphocholine (PC(18:0p/18:1(9Z))), 1-stearoyl-2-hydroxy-sn-glycero-3-phosphocholine (LPC(18:0)), 1-oleoyl-2-hydroxy-sn-glycero-3-phosphocholine (LPC(18:1)), 1-palmitoyl-2-oleoyl-sn-glycero-3-phosphoethanolamine (PE(16:0/18:1)), 1-(1Z-octadecenyl)-2-docosahexaenoyl-sn-glycero-3-phosphocholine (PC(18:0p/22:6)) and 1-stearoyl-2-linoleoyl-sn-glycerol (DG(18:0/18:2)), 1-(9Z-octadecenoyl)-sn-glycero-3-phosphoethanolamine (LPE(18:1)), N-(9Z-octadecenoyl)-sphinganine (Cer(d18:0/18:1(9Z))), 1-hexadecyl-2-(9Z-octadecenoyl)-sn-glycero-3-phosphoethanolamine (PE(16:0/18:1)) from Avanti Polar Lipids, 1-Palmitoyl-2-Hydroxy-sn-Glycero-3-Phosphatidylcholine (LPC(16:0)), 1,2,3 trihexadecanoalglycerol (TG(16:0/16:0/16:0)), 1,2,3-trioctadecanoylglycerol (TG(18:0/18:0/18:0) and 3β-hydroxy-5-cholestene-3-stearate (ChoE(18:0)), 3β-Hydroxy-5-cholestene-3-linoleate (ChoE(18:2)) from Larodan, were prepared to the following concentration levels: 100, 500, 1000, 1500, 2000 and 5000 ng/mL (in CHCl3:MeOH, 2:1, v/v) including 1250 ng/mL of each internal standard. The curves had R values >0.99 for all lipids. Identification was done based on *in-house* library (m/z, MS/MS, retention times) that is based on analysis of authentic standards. Lipid abbreviations are as follows: lipid headgroup followed by the fatty acyl composition (number of carbons in the fatty acyl, number of double bonds). Lipid headgroups: Ceramide (Cer), Sphingomyelin (SM), Phosphatidylcholine (PC), Alkylphosphatidylcholine (PC-O), Phosphatidylcholine plasmalogen (PC-P), Lysophosphatidylcholine (LPC), Phosphatidylethanolamine (PE), Akylphosphatidylethanolamine (PE-O), Phosphatidylethanolamine plasmalogen (PE-P), Lysophosphatidylethanolamine (LPE), Phosphatidylinositol (PI), Phosphatidylserine (PS), Cholesterol Ester (Ce), Diacylglycerol (DG), Triacylglycerol (TG).

Standard solutions extracted blanks (n=3), pooled QC samples (n=3, an aliquot of each sample pooled), in-house serum QC and NIST SRM 1950 (human plasma) were analyzed together with the samples. Identification was done based on in-house library with retention time and spectral data (level 1 identification, based on Metabolic Standard Initiative^19^) and based on MS spectra using on-line spectral libraries (level 2 identification). For pooled samples, identified lipids had an average RSD in the pooled cell samples 15.1% and in extracellular media RSD was on average 16.8%. For polar and semipolar compounds, the RSDs were 25.1% and 26.5% for cells and extracellular media, respectively.

### Data preprocessing

Processing of MS data was performed using the open-source software package MZmine 4.5.0.^20^ The following steps were applied in this processing: (i) Crop filtering with a m/z range of 350 – 1200 m/z and an RT range of 2.0 to 12 minutes, (ii) Mass detection with a noise level of 750 for MS 1 and 200 for MS 2 level, (iii) Chromatogram builder with a minimum time span of 0.08 min, minimum height of 1000 and a m/z tolerance of 0.006 m/z or 10.0 ppm, (iv) Chromatogram deconvolution using the local minimum search algorithm with a 70% chromatographic threshold, 0.05 min minimum RT range, 5% minimum relative height, 1200 minimum absolute height and minimum ration of peak top/edge of 1.2, (v) Isotopic peak grouper with a m/z tolerance of 5.0 ppm, RT tolerance of 0.05 min, maximum charge of 2 and with the most intense isotope set as the representative isotope, (vi) Join aligner with a m/z tolerance of 0.009 or 10.0 ppm and a weight for of 2, a RT tolerance of 0.15 min and a weight of 1 and with no requirement of charge state or ID and no comparison of isotope pattern, (vii) Peak list row filter with a minimum of 10% of the samples (viii) Gap filling using the same RT and m/z range gap filler algorithm with an m/z tolerance of 0.009 m/z or 10.0 ppm, (ix) Identification of lipids and metabolites using a custom database search with an m/z tolerance of 0.007 m/z or 8.0 ppm and a RT tolerance of 0.25 min, and (x) Normalization using internal standards: For lipids: PE(17:0/17:0), SM(d18:1/17:0), Cer(d18:1/17:0), LPC(17:0), TG(17:0/17:0/17:0) and PC(16:0/d30/18:1)) for identified lipids and closest internal standard for the unknown lipids followed by calculation of the concentrations based on lipid-class concentration curves. For polar metabolites the following internal standards were used: valine-d8, glutamic acid-d5, succinic acid-d4, heptadecanoic acid, lactic acid-d3, citric acid-d4. 3-hydroxybutyric acid-d4, arginine-d7, tryptophan-d5, glutamine-d5, CA-d4, CDCA-d4, GCA-d4, GCDCA-d4, GLCA-d4, GUDCA-d4, LCA-d4, TCA-d4, UDCA-d4 and closest internal standard for the unknown metabolites followed by calculation of the concentrations-based concentration curves. For data filtering, we have removed compounds that were present at blank samples (peak area >5 times that of blank) and compounds that had RSD>30% in the pooled quality control samples. MS/MS data was done for the pooled quality control samples using auto MS/MS mode. The two cell lines were analyzed separately but the data preprocessing was done together. The data was adjusted by in-house pooled samples that were analyzed in both batches to correct the difference between the two batches.

### Data analysis

Data analyses were conducted using MetaboAnalyst 6.0^21^ and the R statistical programming language (version 4.1.2) (https://www.r-project.org/). In order to correct for heteroscedasticity, the exposure datasets were pre-processed by log10 transformation and scaling to zero mean and unit variance (autoscaled) as metabolomics data is not normally distributed. The statistical analyses included principal component analysis (PCA), analysis of variance (ANOVA), t-test and fold-change between controls and individual PFAS concentrations, Spearman correlations, and partial correlations between the PFAS and metabolites.

Pathway enrichment analysis was performed using the MetaboAnalyst 6.0 web platform with the Functional Analysis (MS Peaks) module.^22^ This approach supports functional analysis of untargeted metabolomics data generated from high-resolution mass spectrometry. The pathway analysis was done with the data of the polar and semipolar metabolites, as the pathway analysis for lipidomics data is not sufficiently robust due to lack of exact structures of the lipids (fatty acid composition, including the position of the double bonds, cis/trans configuration). However, our polar/semipolar panel (Method 1) includes a large number of lipids, except for neutral lipids (CE, DG, TG) that are not covered either by sample preparation nor the negative ion mode. The input data for the pathway analysis comprised complete LC-HRMS data, i.e. both identified and unknown metabolites, obtained in negative ionization mode. First, we performed statistical analyses using t-test between control and each exposure concentration between PFAS and polar metabolites, resulting in fold change, p values and FDR values. The whole input peak list, with peak names given as their numeric mass (m/z) values for putative annotation, and the statistical results with False Discovery Rate (FDR)-corrected p-values and t-score was used for the pathway analysis. Two algorithms were applied separately, namely, Mummichog and Gene Set Enrichment Analysis (GSEA) and two pathway libraries were used in the pathway analysis, namely human scale metabolic model MNF (from MetaboAnalyst Mummichog package) and Kyoto Encyclopedia of Genes and Genomes (KEGG) pathways for *Homo Sapiens* to determine the relative significance of the identified pathways (Li et al. 2020). The mass tolerance for the pathway analysis was set at 7 ppm, and we also used an advanced option to select representative adducts by removing isotopic adducts as these have been already removed in our data preprocessing step.

## Results

### Global metabolic responses to perfluorohexyloctane exposure

To evaluate the metabolic effects of F6H8, we performed untargeted metabolomic including lipidomic profiling of hepatocytes and their extracellular environment across a wide range of non-cytotoxic exposure concentrations.

Several compounds, both in the cells as well as in the extracellular media, showed a strong linear relationship with F6H8 concentration (**Supplementary Table 1**). Notably, three unknown compounds in both cells and media showed a very strong (R>0.78, FDR < 1.9e^-16^) correlation with the exposure concentration, and these compounds were not detected in control cells nor in control media. For two of the compounds, we could obtain mass spectra. Based on MS/MS spectra, one of these compounds could be a derivative of F6H8, *i.e.*, F6H8 biotransformation product. The compound was identified as a carboxylic-acid derivative of F6H8, namely perfluorohexyloctanoic acid (Figure 1A). The deprotonated molecular ion, [M–H]⁻, matched that of perfluorohexyloctanoic acid with a mass error < 1 ppm. In the negative-ion MS/MS spectrum, the product ions arise from neutral loss of the carboxyl group and from sequential, extensive HF eliminations. This loss pattern has previously been observed both on H-substituted PFAS (*e.g.*, fluorotelomer carboxylic acids)^23, 24^ and semi-fluorinated alkanes.^25^ In addition to the mass spectral data, also the retention time (rt) of the compound (5.67 min) was in line with the corresponding fully fluorinated carboxylic acid (Perfluoro-n-tetradecanoic acid, rt = 5.74 min) which had a slightly higher retention time than partially fluorinated compound. We also performed a retention time prediction, based on the logP value of perfluorohexyloctanoic acid, computed to be 6.9 according to Pubchem model XlogP3 3.0 (**Supplementary Figure 3**). Using the linear regression equation, the predicted retention time of the compound is 5.77 min, which is very close to the retention time measured, with Δrt = 0.07 min. The second unknown compound of interest showed a typical fragment of sulfonic group (**Figure 1B**), and co-clustered with other sulphate containing metabolites in ion network analysis. However, majority of the compounds that correlated with the F6H8 concentration could not be identified.

**Figure 1.**
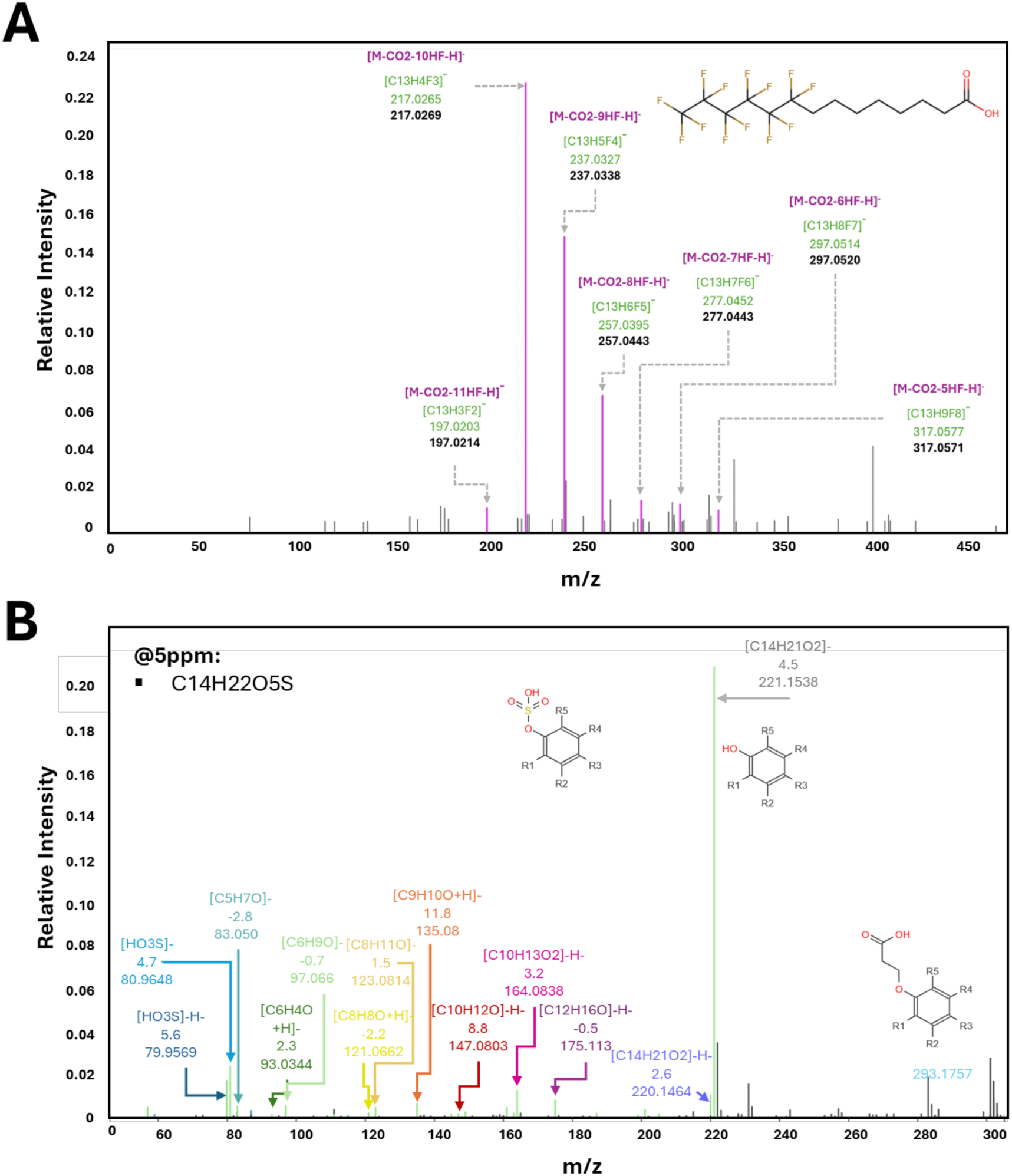
**(A)** MS/MS spectrum of Compound 203 (m/z = 4C1.0781, rt 5.C7 min) in cells was linearly correlating with the FCH8 exposure concentration (R=0.78, p=1.81e^-18^, FDR=1.S4e^-1C^). (**B)** MS/MS spectrum of Compound 203 CC (m/z = 301.1103, rt 2.S8 min) in cells was linearly correlating with the FCH8 exposure concentration (R=0.8S, p=1.5S^-2S^, FDR=3.4e^-27^).

Overall, 65 polar/semipolar compounds and 66 lipids showed significant correlation (FDR < 0.05) with F6H8 concentration, among which 23 were negatively and the remainder positively correlated. Of the identified compounds, several free fatty acids (FFAs) were negatively associated with the exposure as well as one amino acid, one acylcarnitine, and two lipids. Large number of lipids, amino acids, two bile acids, and some sugar derivatives were positively correlated with the F6H8 concentration.

### Lipidome alterations in hepatocytes and extracellular media

Exposure to F6H8 caused concentration-dependent, non-monotonic changes in lipid profiles in both cells and extracellular media, as indicated by fold changes relative to the controls (**Figure 2, Supplementary Tables 2-3**). The exposure caused a clear increase of lipids in the extracellular media, except for hexadecenoylcarnitine, which was downregulated. Particularly phospholipids and several triacylglycerols (TGs) were upregulated in the media, and both lysophospholipids and PC(35:4) exhibited increasing upregulation with higher F6H8 concentrations. We also observed dose-dependent increase in polyunsaturated FFAs, while the saturated FFAs were downregulated with increasing F6H8 exposure. In addition, several amino acids and their derivatives were downregulated in response to F6H8 exposure.

**Figure 2.**
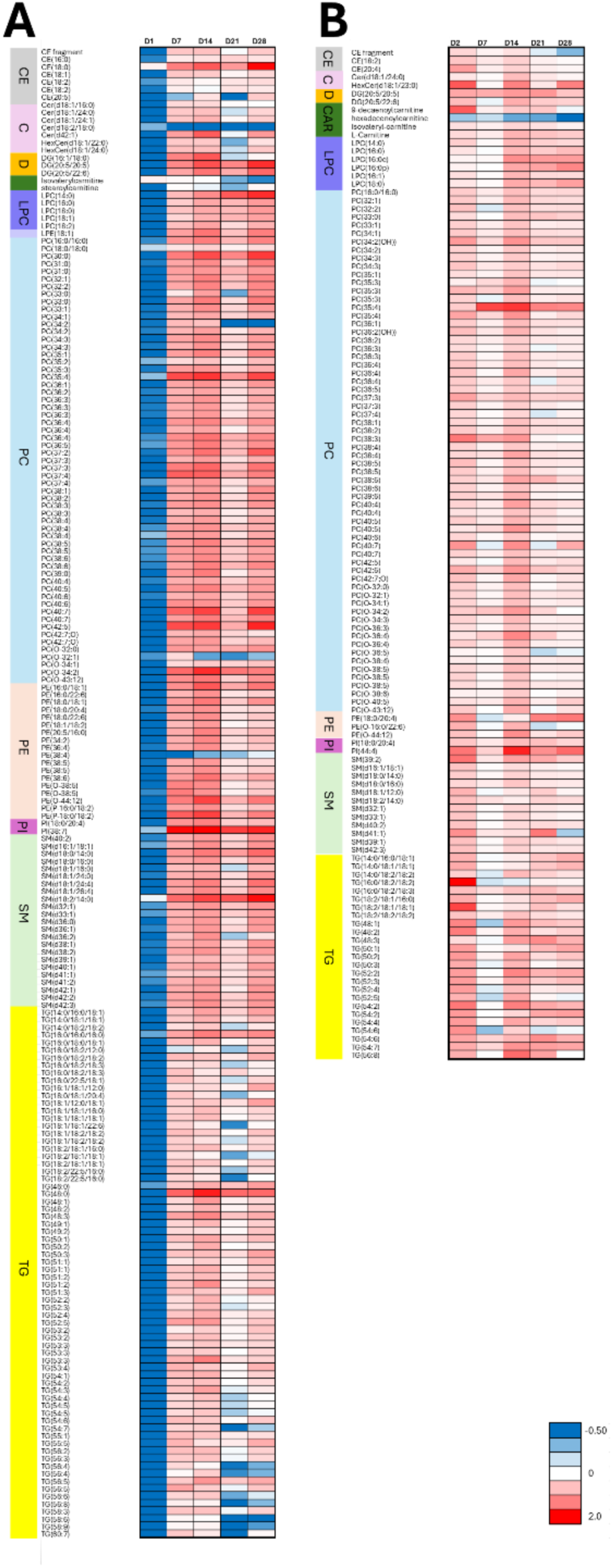
Fold changes in lipids levels between F6H8-exposed and control HepaRG cells across different exposure concentrations, shown for both **(A)** cells and **(B)** extracellular media. Only those fold changes that are significant at least in one exposure concentration are shown. Lipid class abbreviations: CE – cholesteryl ester, C – ceramide, LPC – lysophosphatidylcholine, PC – phosphatidylcholine, PI – phosphatidyl inositol, PE – phosphatidylethanolamine, SM – sphingomyelin, TG – triacylglycerol, DG – diacylglycerol.

In cells, there was even more substantial effect on lipid profiles, with the lowest F6H8 concentration causing marked reduction of large number of lipids, with inverse trend at the higher exposures, with several lipids upregulated following exposure. For example, ceramide Cer(d18:2/18:0) was consistently downregulated at all tested concentrations, while sphingomyelin SM(d18:2/14:0) was upregulated at the higher exposure levels.

Given the widespread lipid alterations, we also investigated the impact of exposure on the level of lipid classes and lipid class ratios. Lysophosphatidylcholines (LPCs) and LPC/PC (PC – phosphatidylcholine) ratio had a significant positive correlation with the exposure concentration. We also performed partial correlation analysis across the exposure groups for the lipid classes, comparing the correlation patterns in control cells and exposed cells (**Figure 3**). Marked positive correlations observed between different lipid classes in control cells were altered following the exposure to F6H8, with less intra-class positive correlations and strong negative correlations between polyunsaturated fatty acid (PUFA)-containing TGs and (i) LPC/PC and PC_PUFA at the lowest exposure concentration. and (ii) monounsaturated fatty acids (MUFA) containing TG and saturated fatty acids (SFA) containing PCs at the highest F6H8 concentration. Also, the association between phosphatidylethanolamines (PEs) and SMs was lost after the F6H8 exposure.

**Figure 3.**
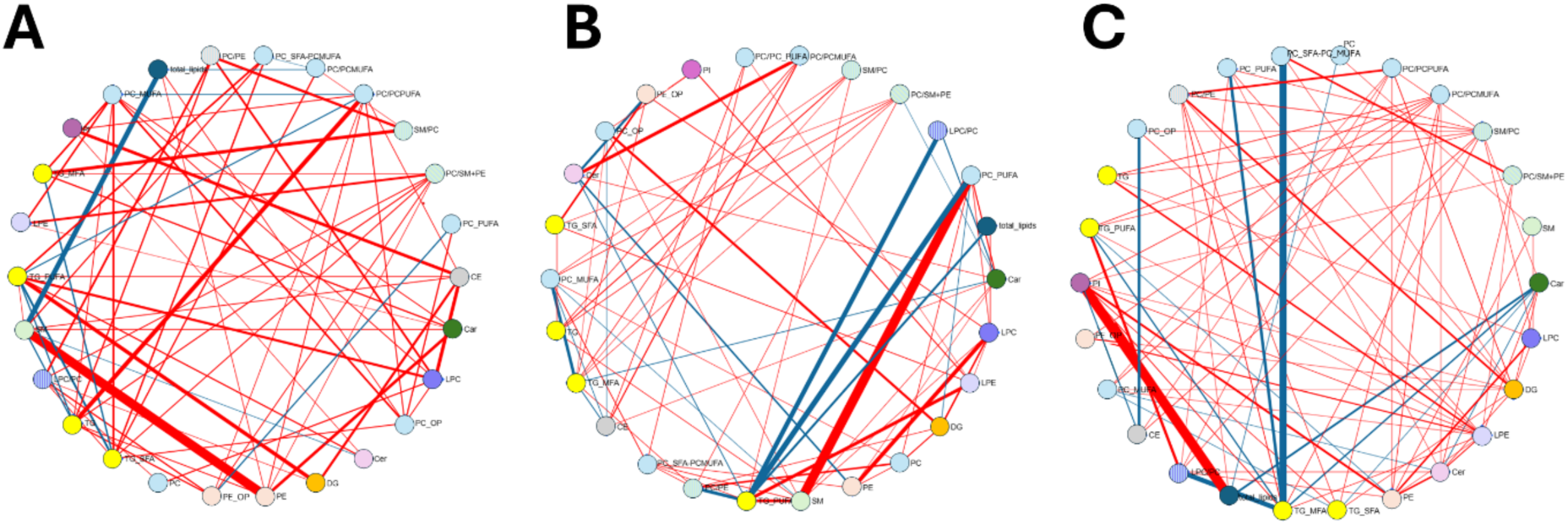
Partial correlation networks including lipid classes as node, in **(A)** control HepaRG cells, **(B)** cells exposed to the lowest F6H8 concentration, and **(C)** cells exposed to the highest FCH8 concentration. The networks illustrate reduced interclass connectivity between lipid species following exposure. The thickness of lines represents the strength of correlation, with red lines representing positive correlation, and blue lines negative correlation.

Lipid class abbreviations: CE – cholesteryl ester, LPC – lysophosphatidylcholine, PC – phosphatidylcholine, PI – phosphatidyl inositol, PE – phosphatidylethanolamine, SM – sphingomyelin, TG – triacylglycerol, DG – diacylglycerol. For phospholipids, OP states alkylether derivatives of parent phospholipid, for TGs and phospholipids, SFA defines saturated fatty acyls, MUFA monounsaturated fatty acyls and PUFA polyunsaturated fatty acyls.

### Effects on polar and semipolar metabolite profiles

We also investigated the impact of exposure on polar and semipolar metabolites. Similar to lipids, we observed concentration-dependent, non-monotonic changes both in cells and extracellular media (**Figure 4, Supplementary Tables 2-3**). In cells, four amino acids (5-oxoproline, aspartic acid, ketoleucine, and threonine) were significantly altered, with varying pattern of changes. Aspartic acid and ketoleucine were downregulated at higher F6H8 exposure concentrations. Among FFAs, three were significantly affected, with eicosatetraenoic acid showing a linear decrease with the increasing F6H8 concentration. In the extracellular media, FFAs showed patterns that were linked with their saturation level: saturated FFAs were downregulated while mviabioityonounsaturated (MUFA) and polyunsaturated (PUFA) fatty acids were upregulated, particularly at the highest concentration. Amino acids and their derivatives generally exhibited a downregulation trend.

**Figure 4.**
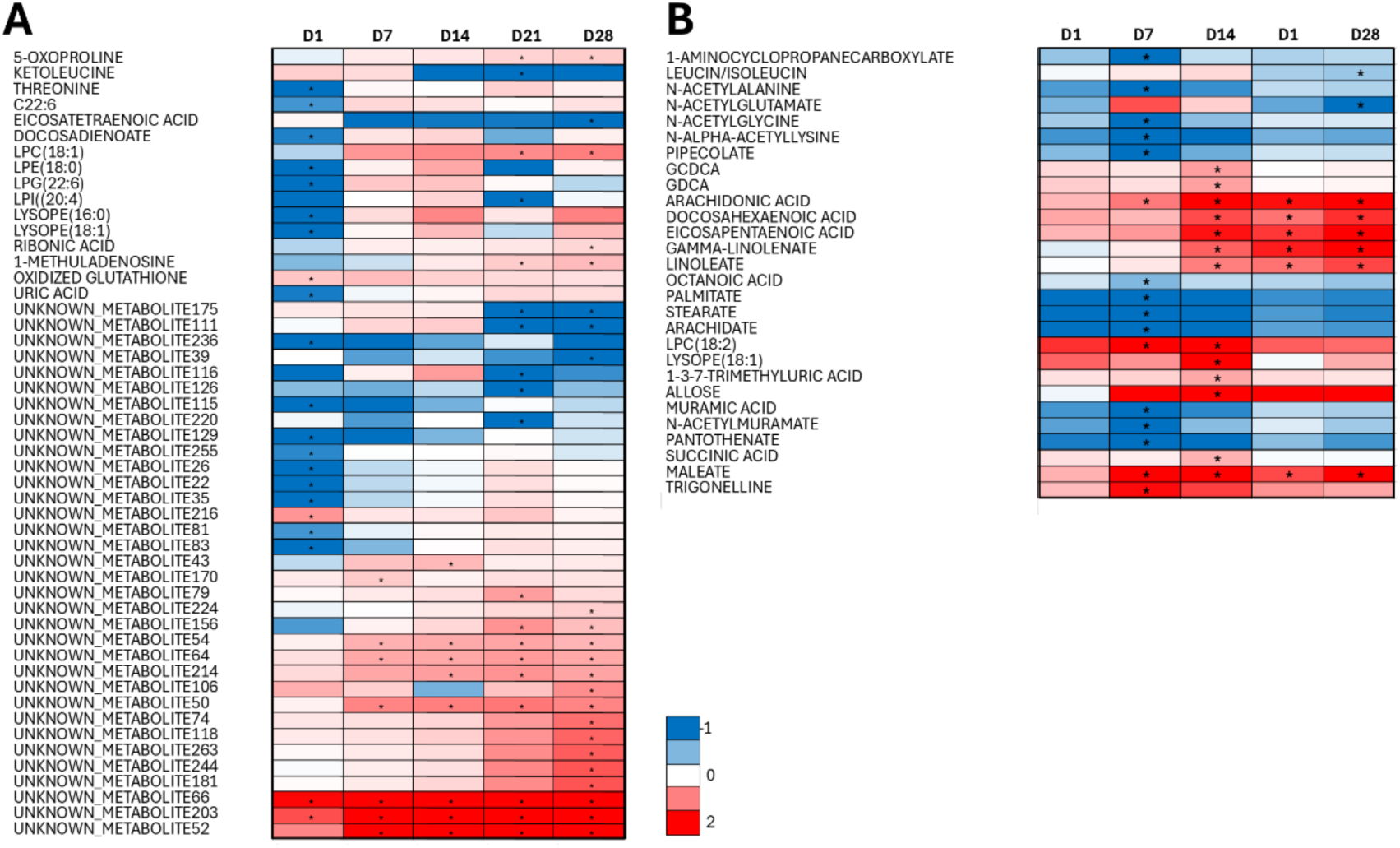
Fold changes in polar/semipolar metabolites between F6H8-exposed samples and control cells and across different exposure concentrations in **(A)** HepaRG cells, and **(B)** in extracellular media. Only those fold changes that are significant at least in one exposure level are shown. For extracellular media, only identified compounds are shown. For detailed information, see **Supplementary Table 3**.

### Pathway enrichment and functional analysis

We also performed both lipid enrichment and functional analysis at the pathway level for the lowest and highest F6H8 concentrations, which showed opposite metabolic patterns (**Table 2**). Four pathways were affected at both concentrations (Biosynthesis of unsaturated fatty acids, Fatty Acid Metabolism, Chondroitin sulfate degradation and Heparan sulfate degradation) but showed opposite trends: all were downregulated at the lowest concentration and upregulated at the highest. At the lowest F6H8 concentration, a large number of pathways related to fatty acid and lipid metabolism were altered as well as pathways related to amino acid and nitrogen metabolism. Majority of the affected pathways were downregulated, only Keratan sulfate biosynthesis was upregulated at the lowest exposure concentration. Notably, sphingolipid metabolism was among the most prominently affected pathways. At the highest F6H8 concentration, alterations were observed in several amino acids and nitrogen metabolism-related pathways, along with some fatty acid related pathways. In addition, the metabolic pathways linked with central carbon metabolism were also affected. Central carbon and energy metabolism related pathways were mainly downregulated while amino acids and nitrogen metabolism-related pathways showed alteration in both directions, indicating complex regulatory responses at higher exposure levels.

**Table 2.** Summary of pathway enrichment analysis for the lowest (ED1) and highest (ED28) F6H8 exposure concentrations. Pathways showing significant enrichment (p < 0.05) are listed along with their direction of change.

| Comment | Pathway | ED1 | p ED1 | ED28 | p ED28 |
| --- | --- | --- | --- | --- | --- |
| Amino Acid & Nitrogen Metabolism | Beta-alanine degradation I | ns |  | up | 0.011 |
| Amino Acid & Nitrogen Metabolism | Aminosugars metabolism | down | 0.039 | ns |  |
| Amino Acid & Nitrogen Metabolism | Arginine and proline metabolism | ns |  | up | 0.028 |
| Amino Acid & Nitrogen Metabolism | Citrulline biosynthesis | down | 0.022 | ns |  |
| Amino Acid & Nitrogen Metabolism | Glutamate metabolism | ns |  | down | 0.042 |
| Amino Acid & Nitrogen Metabolism | Glycine, serine and threonine metabolism | ns |  | down | 0.031 |
| Amino Acid & Nitrogen Metabolism | Methionine and cysteine metabolism | ns |  | down | 0.049 |
| Amino Acid & Nitrogen Metabolism | proline biosynthesis II (from arginine) | ns |  | up | 0.009 |
| Amino Acid & Nitrogen Metabolism | threonine degradation II | down | 0.004 | ns |  |
| Amino Acid & Nitrogen Metabolism | Urea cycle/amino group metabolism | ns |  | up | 0.044 |
| Central carbon and energy metabolism | Butanoate metabolism | ns |  | up | 0.029 |
| Central carbon and energy metabolism | Citrate cycle (TCA cycle) | ns |  | down | 0.029 |
| Central carbon and energy metabolism | D-glucuronate degradation I | ns |  | up | 0.009 |
| Central carbon and energy metabolism | Glyoxylate and dicarboxylate metabolism | ns |  | down | 0.024 |
| Central carbon and energy metabolism | Propanoate metabolism | ns |  | down | 0.025 |
| Central carbon and energy metabolism | Pyruvate metabolism | ns |  | down | 0.038 |
| Lipid and fatty acid metabolism | Biosynthesis of unsaturated fatty acids | down | 0.008 | up | 0.033 |
| Lipid and fatty acid metabolism | De novo fatty acid biosynthesis | down | 0.021 | ns |  |
| Lipid and fatty acid metabolism | Fatty acid activation | down | 0.029 | ns |  |
| Lipid and fatty acid metabolism | Fatty Acid Metabolism | down | 0.050 | up | 0.045 |
| Lipid and fatty acid metabolism | Glycerolipid metabolism | ns |  | up | 0.019 |
| Lipid and fatty acid metabolism | Glycosphingolipid biosynthesis - ganglioseries | down | 0.016 | ns |  |
| Lipid and fatty acid metabolism | Glycosphingolipid biosynthesis - globoseries | down | 0.013 | ns |  |
| Lipid and fatty acid metabolism | Glycosphingolipid biosynthesis - lactoseries | down | 0.031 | ns |  |
| Lipid and fatty acid metabolism | Glycosphingolipid biosynthesis – neolactoseries | down | 0.045 | ns |  |
| Lipid and fatty acid metabolism | Glycosphingolipid metabolism | down | 0.042 | ns |  |
| Lipid and fatty acid metabolism | leukotriene biosynthesis | down | 0.004 | ns |  |
| Lipid and fatty acid metabolism | Leukotriene metabolism | down | 0.024 | ns |  |
| Lipid and fatty acid metabolism | Linoleic acid metabolism | down | 0.023 | ns |  |
| Lipid and fatty acid metabolism | Phosphatidylinositol phosphate metabolism | down | 0.013 | ns |  |
| Lipid and fatty acid metabolism | Phytanic acid peroxisomal oxidation | ns |  | down | 0.038 |
| Glycosaminoglycan turnover | Chondroitin sulfate degradation | down | 0.030 | up | 0.038 |
| Glycosaminoglycan turnover | Heparan sulfate degradation | down | 0.030 | up | 0.038 |
| Glycosaminoglycan turnover | Hyaluronan Metabolism | down | 0.017 | ns |  |
| Glycosaminoglycan turnover | Keratan sulfate biosynthesis | up | 0.031 | ns |  |
| Glycosaminoglycan turnover | Keratan sulfate degradation | down | 0.030 | ns |  |
| Protein glycosylation | N-Glycan biosynthesis | down | 0.019 | ns |  |
| Protein glycosylation | O-Glycan biosynthesis | down | 0.045 | ns |  |
| Cofactor and Vitamin Metabolism | Porphyrin metabolism | down | 0.039 | ns |  |
| Cofactor and Vitamin Metabolism | Pyrimidine metabolism | down | 0.025 | ns |  |
| Nucleotide and Translation Machinery | tRNA charging | down | 0.026 | ns |  |
| Nucleotide and Translation Machinery | Vitamin B6 (pyridoxine) metabolism | down | 0.045 | ns |  |

## Discussion

In this study, we observed mass spectral features consistent with fluorinated metabolites in F6H8-treated hepatocytes. One of these was tentatively annotated as a carboxylic acid derivative of F6H8, tridecafluorotetradecanoic acid, suggesting potential oxidative biotransformation. This observation aligns with previous studies showing that structurally analogous non-fluorinated saturated alkanes can undergo hepatic ω-oxidation to yield carboxylic acids of the same chain length.^10–12^ The proposed alkane oxidation pathway in hepatocytes involves sequential cytochrome P450-mediated hydroxylation of the terminal methyl group to a primary alcohol, followed by oxidation to an aldehyde *via* alcohol dehydrogenase and subsequent conversion to the corresponding carboxylic acid by aldehyde dehydrogenase. Given the structural similarity to alkanes, F6H8 may undergo biotransformation through an analogous pathway.

Although definitive structural confirmation was not possible due to the absence of analytical reference standards, the detection of fluorine-containing features supports the possibility of intracellular metabolism of F6H8. This warrants further investigation, as such carboxylic acid metabolites could possess greater bioactivity, persistence, or toxicity than the parent compound.

We observed that hepatocytes actively secreted lipids following exposure to F6H8 across all tested concentrations, as evidenced by elevated lipid levels in the extracellular medium. This response may reflect F6H8-induced metabolic dysregulation, or indicate membrane perturbations that trigger remodelling, breakdown, or stress responses such as endoplasmic reticulum (ER) stress or oxidative stress. One plausible explanation for the increased lipid efflux is the ability of F6H8 to associate with and permeate cellular lipid membranes. Semifluorinated alkanes have been reported to co-disperse with phospholipids in bilayers, where their lipophobic fluorinated chains segregate to form a phase-separated fluorinated core within the membrane.^14^ Such reorganization could alter membrane fluidity and facilitate dysregulated lipid trafficking .

Notably, lysophospholipids (LPLs) exhibited a concentration-dependent and non-monotonic pattern: intracellular levels decreased at low concentrations but increased markedly at higher exposures in both cells and extracellular media. This accumulation may result from enhanced LPL synthesis or impaired degradation under high-exposure conditions. Concurrently, the consistent downregulation of acylcarnitines in both compartments suggests suppression of mitochondrial β-oxidation, potentially compromising energy metabolism and fatty acid homeostasis.

F6H8 exposure was associated with concentration-dependent alterations in multiple metabolic pathways, many of which displayed non-monotonic responses. At the lowest concentration, pathway analysis revealed signatures of oxidative stress and inflammatory activation, including perturbations in leukotriene, linoleic acid, phosphoinositide (PIP), glycosphingolipid, and aminosugar metabolism. Dysregulation of several sphingolipid-related pathways further supports a hepatocellular stress response. Sphingolipids serve as key signaling molecules regulating cell proliferation, adhesion, migration, autophagy, apoptosis, and mitochondrial function.^26^ Accordingly, disruptions in these pathways may reflect adverse impacts on core cellular processes. Their suppression suggests broad lipid metabolic reprogramming—potentially an adaptive response to metabolic stress or toxicity. Moreover, altered sphingolipid metabolism has been implicated in type 2 diabetes, metabolic syndrome, and drug-induced liver injury,^27, 28^ implying that F6H8 may influence pathways relevant to these conditions.

Interestingly, several pathway-level similarities were observed between the current findings and our previous study on PFAS-exposed hepatocytes,^29^ particularly at the lowest exposure concentration, where both exposures affected fatty acid activation, *de novo* fatty acid biosynthesis, PIP metabolism, glycosphingolipid metabolism, pyrimidine metabolism, and O-glycan biosynthesis. In contrast, fewer overlaps were evident at the highest F6H8 concentration, with only β-alanine metabolism, butanoate metabolism, and urea/amino group metabolism being commonly affected. At the highest exposure levels, pronounced disruptions were observed in central carbon metabolism, including changes in tricarboxylic acid (TCA) cycle intermediates and amino acid degradation pathways (notably β-alanine, glutamate, lysine, and isoleucine). These alterations point to mitochondrial dysfunction, impaired nutrient-sensing pathways, and elevated oxidative stress.

The non-monotonic nature of the response aligns with previous findings for conventional PFAS, observed both in hepatocyte models^30^ and in human occupational exposure studies.^31^ However, the underlying metabolic fingerprints differ. PFAS exposure typically results in downregulation of bile acids, acylcarnitines, and free fatty acids, whereas F6H8 exposure primarily affected lipid and amino acid metabolism, with minimal effects on bile acids. While PFAS are known to impair bile acid synthesis *via* CYP7A1 inhibition,^32^ it is unlikely that F6H8 acts through similar receptor-mediated mechanisms given its chemical structure. Instead, F6H8 may elicit its effects through pathways shared with structurally related alkanes, which have been shown to induce lipid peroxidation, increase membrane permeability, and reduce cell viability. For instance, alkanes such as nonane (C9), decane (C10), undecane (C11), and dodecane (C12) decrease cell viability in *Saccharomyces cerevisiae* by disrupting membrane integrity.^33^ These findings suggest that F6H8 could similarly accumulate within hepatocytes and interfere with membrane-associated processes, leading to metabolic disturbances.

Overall, the non-monotonic metabolic changes observed in response to F6H8 indicate a biphasic response, with adaptive mechanisms activated at low concentrations that become overwhelmed or dysregulated at higher exposures. Such changes may also result from the formation of F6H8 metabolites that are more bioactive or toxic than the parent compound. The consistent elevation of extracellular lipid levels across all concentrations further supports the hypothesis of membrane leakage or enhanced secretion as a manifestation of membrane and metabolic stress.

Several limitations should be acknowledged. Exposure levels were estimated based on data from structurally related compounds due to the absence of pharmacokinetic data on F6H8 absorption, distribution, and tissue accumulation. A previous study in rabbits^8^ detected F6H8 in blood at high parts-per-billion (ppb) concentrations and indicated that repeated administration could lead to systemic enrichment. Given the lipophilic nature of F6H8 and evidence from similar compounds, hepatic accumulation is plausible. Additionally, while our exposure duration followed established hepatocyte model protocols, it may not fully represent chronic exposure scenarios. Nevertheless, this model has been validated in prior studies investigating PFAS mixtures, producing metabolic patterns consistent with both *in vivo* animal data and human exposure profiles.^30^

Taken together, this study provides the first comprehensive evaluation of the metabolic effects of perfluorohexyloctane (F6H8) in human hepatocytes. We identified several putative biotransformation products, suggesting that F6H8 may undergo metabolic conversion despite prior assumption of inertness. Exposure induced pronounced, concentration-dependent disruptions in lipid and energy metabolism, accompanied by molecular signatures indicative of oxidative stress, mitochondrial dysfunction, and membrane perturbations. These findings raise concerns about the systemic safety of F6H8 in medical applications and highlight the need for further investigation into its biological activity and long-term effects. Although limited by the *in vitro* design and estimated exposure parameters, the observed non-monotonic and complex metabolic responses underscore the importance of comprehensive toxicological assessment of F6H8 and related semifluorinated alkanes.

## Acknowledgments

This study was supported by the Swedish Research Council (grants no. and 2020-03674 and 2016-05176 to T.H and M.O), Formas (grant no. 2019-00869 to T.H and M.O), Novo Nordisk Foundation (Grants no. NNF20OC0063971 and NNF21OC0070309 to T.H. and M.O.), and by the “Investigation of endocrine-disrupting chemicals as contributors to progression of metabolic dysfunction-associated steatotic liver disease” (EDC-MASLD) consortium funded by the Horizon Europe Program of the European Union under Grant Agreement 101136259 (to MO and TH). The study was also partially supported by grants from the Swedish Knowledge Foundation (Grants. no. 20160019; 20220122). We also thank Lorane Valgueblasse for help in the sample preparations.

## Disclaimer

Views and opinions expressed are those of the author(s) only and do not necessarily reflect those of the European Union. Neither the European Union nor the granting authority can be held responsible for them.

## Author contributions

Andi Alijagic: Methodology, Validation, Formal analysis, Investigation, Resources, Writing - Original Draft, Writing - Review & Editing, Visualization, Project administration, Funding acquisition.

Jade Chaker: Methodology, Formal analysis, Visualization, Writing - Review C Editing.

Joao Barbosa: Methodology, Formal analysis, Writing - Review C Editing.

Daniel Duberg: Methodology, Writing - Review C Editing

Victor Castro Alves: Methodology, Formal analysis, Writing - Review C Editing.

Alex Dickens: Methodology, Formal analysis, Writing - Review C Editing.

Matej Orešič: Conceptualization, Writing - Review & Editing, Funding acquisition.

Tuulia Hyötyläinen: Conceptualization, Validation, Formal analysis, Investigation, Resources, Writing - Original Draft, Writing - Review & Editing, Visualization, Project administration

