## Supplementary Figures for "Metabolic effects and biotransformation of perfluorohexyloctane (F6H8) in human hepatocytes"

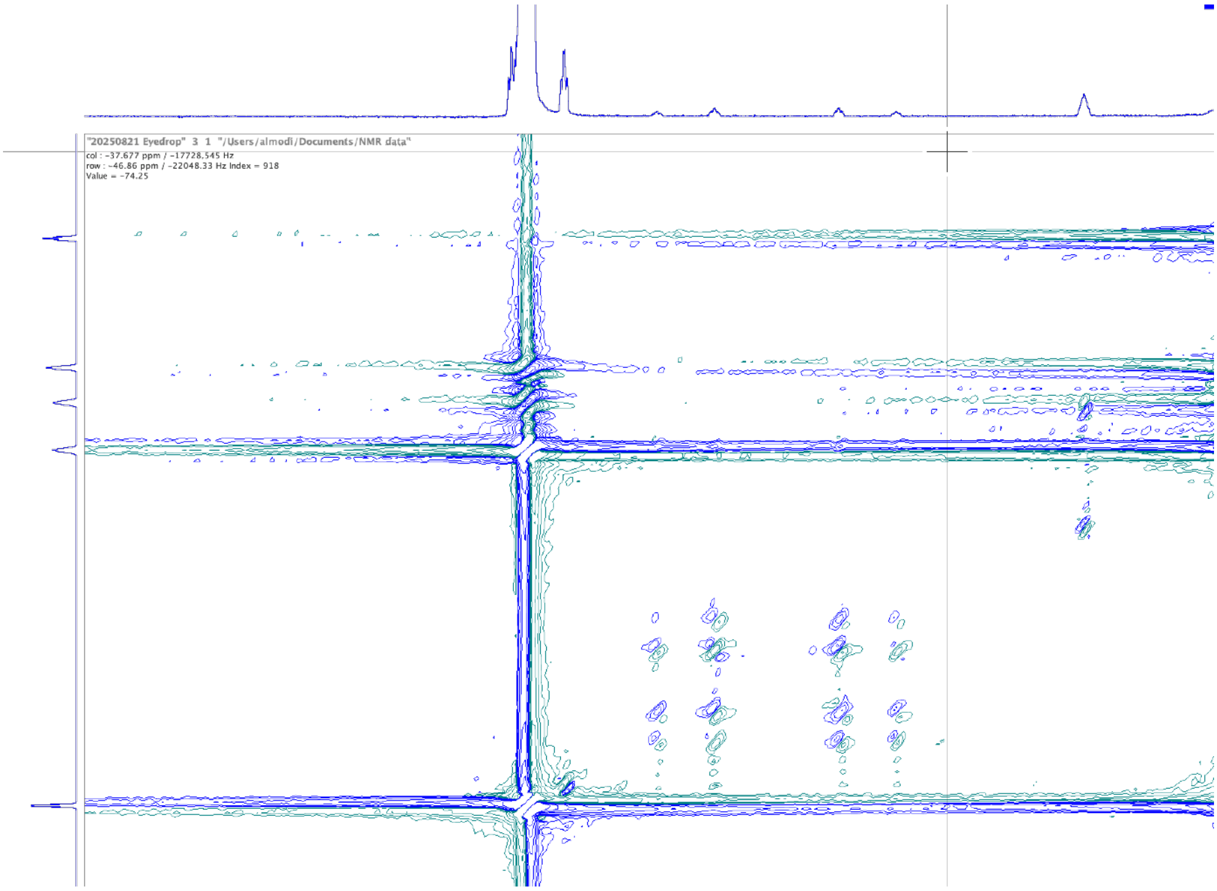


**Supplementary Figure 1.** NMR spectrum of the eyedrop solution, showing some potential fluorinated impurities.

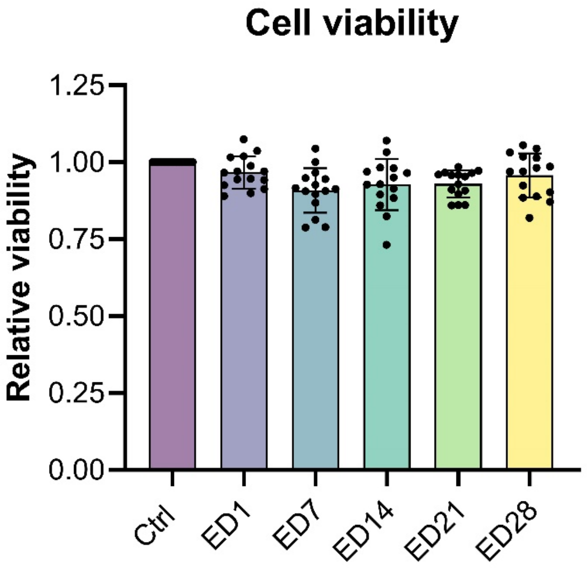


**Supplementary Figure 2.** Results of the alamarBlue viability assay assessing the effects of perfluorohexyloctane exposure on HepaRG cells. Bars represent the mean and standard deviation of three independent biological replicates, each consisting of five technical replicates per exposure concentration. Viability is expressed as relative fold change compared to the negative (unexposed) control.

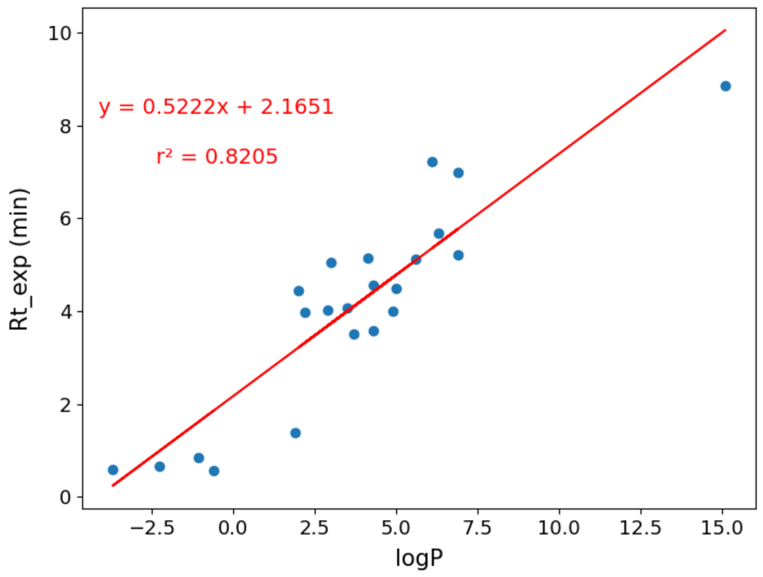


**Supplementary Figure 3.** Linear regression of experimental retention time (RT_exp) of internal standards against their octanol-water coefficient (logP).
